# Cell-surface receptors enable perception of extracellular cytokinins

**DOI:** 10.1101/726125

**Authors:** Ioanna Antoniadi, Ondřej Novák, Zuzana Gelová, Alexander Johnson, Ondřej Plíhal, Thomas Vain, Radim Simerský, Václav Mik, Michal Karady, Markéta Pernisová, Lenka Plačková, Korawit Opassathian, Jan Hejátko, Stéphanie Robert, Jiří Friml, Karel Doležal, Karin Ljung, Colin Turnbull

**Affiliations:** Department of Life Sciences, Imperial College London, London SW7 2AZ, United Kingdom; Umeå Plant Science Centre, Department of Forest Genetics and Plant Physiology, Swedish University of Agricultural Sciences, SE-901 83 Umeå, Sweden; Laboratory of Growth Regulators, Institute of Experimental Botany of the Czech Academy of Sciences and Faculty of Science of Palacký University, Šlechtitelů 27, CZ-78371 Olomouc, Czech Republic; Institute of Science and Technology Austria, 3400 Klosterneuburg, Austria; CEITEC – Central European Institute of Technology and NCBR, Faculty of Science, Masaryk University, Kamenice 5, CZ-62500 Brno, Czech Republic; Department of Molecular Biology, Centre of the Region Haná for Biotechnological and Agricultural Research, Faculty of Science of Palacký University, Šlechtitelů 27, CZ-78371 Olomouc, Czech Republic; Department of Protein Biochemistry and Proteomics, Centre of the Region Haná for Biotechnological and Agricultural Research, Faculty of Science of Palacký University, Šlechtitelů 27, CZ-78371 Olomouc, Czech Republic; Department of Chemical Biology and Genetics, Centre of the Region Haná for Biotechnological and Agricultural Research, Faculty of Science of Palacký University, Šlechtitelů 27, CZ-78371 Olomouc, Czech Republic; Development, UMR232/DIADE, Institut de Recherche pour le Développement (IRD), Université de Montpellier, 34394 Montpellier, France

## Abstract

Cytokinins are mobile multifunctional plant hormones with roles in development and stress resilience ^1,2^. Although cytokinin receptors are substantially localised to the endoplasmic reticulum ^3–5^, the cellular sites of cytokinin perception continue to be debated ^1,6,7^. Several cytokinin types display bioactivity ^8,9^ and recently a cell-specific cytokinin gradient was reported in roots ^10^. Yet, the importance of spatially heterogeneous cytokinin distribution and the specific cytokinin(s) that account for the different responses remain unclear. Here we show that cytokinin perception by plasma membrane receptors is an effective path for cytokinin response in root cells. Readout from a Two Component Signalling cytokinin-specific reporter (*TCSn::GFP;*^11^) is closely matched to intracellular cytokinin content, yet a proportion of bioactive cytokinins are detected in the extracellular fluid. Using cytokinins covalently linked to beads that could not pass the plasma membrane, we demonstrate that strong *TCSn* activation still occurs and that this response is greatly diminished in cytokinin receptor mutants. Although intracellular receptors play significant roles, we argue for a revision of concepts of cytokinin perception to include the spatial dimensions. In particular, selective ligand-receptor affinities, cellular localisation and tissue distribution of bioactive cytokinins, their receptors, transporters and inactivation enzymes appear all to be components of the signalling regulatory mechanisms.

---

Cytokinins are key hormones regulating cell division and differentiation, root and shoot architecture, senescence and responses to environmental stresses^1^. The active forms are the cytokinin free bases, which comprise a range of *N*^6^-modified adenine molecules^8,9^, especially trans-zeatin (tZ) and isopentenyl adenine (iP). Homeostatic regulation of active cytokinin pools occurs at the level of biosynthesis, and also through metabolic deactivation by glucosylation, phosphoribosylation or irreversible degradation by cytokinin dehydrogenase (CKX)^9,12^. Cytokinin signalling commences with perception of bioactive molecules by hybrid histidine kinases (HKs)^1,13,14^. Several reports show GFP-fused Arabidopsis HKs (AHKs) mainly localised to the endoplasmic reticulum (ER) membrane^3–5^, yet the originally proposed extracellular site of cytokinin perception at plasma membrane receptors^13,15,16^ has never been discounted. Notably, several classes of cytokinin transporters facilitate movement of cytokinins in and out of the cell^7^. Cytokinins binding to receptors trigger a phosphorelay cascade, resulting in activation of B-type ARABIDOPSIS RESPONSE REGULATORS (ARR-B) transcription factors^17^. The cytokinin-responsive synthetic promoter fusion *TCSn::GFP* reflects global *ARR-B* transcriptional activity^11^ and has facilitated *in vivo* monitoring of cytokinin responses, leading to new discoveries about cytokinin function^18,19^. However, it is unclear whether *TCSn::GFP* signal strength is quantitatively related to cellular cytokinin content and uncertainty remains about which active cytokinin(s) are responsible for different responses in the root tip and other tissues.

Analysis of cytokinins in cell protoplasts isolated from *TCSn::GFP* roots using Fluorescence Activated Cell Sorting (FACS) (Fig. 1a) revealed that total cytokinin content was almost three times higher in the cytokinin responsive (GFP+) cells (Fig. 1b). These results are in accordance with recent evidence for a cytokinin gradient within the root tip^10^ that matches the *TCSn::GFP* expression pattern^11^. We further showed a positive correlation between *TCSn::GFP* signal strength and cytokinin content within GFP+ cell populations displaying higher and lower fluorescence (Fig. 1c and Supplementary Fig. 1). Active cytokinins and their riboside precursors were generally enriched in the most fluorescent (GFP^+^_max_) cells, whereas inactive cytokinin glucosides were more equally distributed between the two sub-populations (Fig. 1d). We conclude that increases in *TCSn::GFP* readout, designed to approximate global *ARR-B* transcript levels, are associated with elevated active cytokinin content as the input signal.

**Fig. 1.**
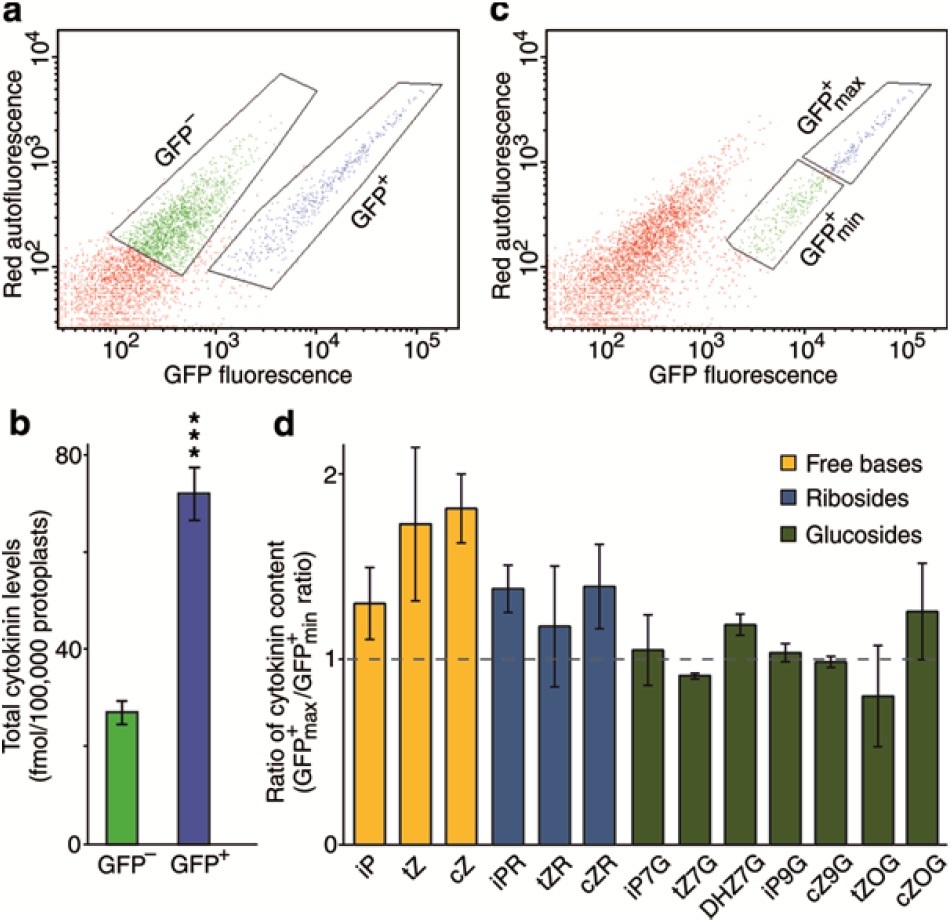
Cell-level cytokinin content correlates with expression of the *TCSn::GFP* cytokinin reporter gene. Analysed by Fluorescence-Activated Cell Sorting (FACS) and liquid chromatography-mass spectrometry (LC-MS). **a**, Autofluorescence scattering intensity plotted against GFP fluorescence intensity for 50,000 events analysed, separating the *TCSn::GFP* root protoplasts into two populations, GFP^−^ cells (green) and GFP^+^ cells (blue) which were then selected (gated) for cell sorting. **b**, Sum of total cytokinin metabolites in sorted GFP^+^ and GFP^−^ cells. Paired sample t-test applied (***, *P* <0.001, n=9 biological replicates; 2 technical replicates for each biological replicate). Error bars are s.e.m. **c**, As for a, but showing selection of *TCSn::GFP*^+^ cell sub-populations with maximum fluorescence (GFP+_max_; blue) and minimum fluorescence (GFP+_min_; green). **d**, Ratios of the concentration of cytokinin metabolites between *GFP*^+^_max_ and GFP^+^_min_ cells (n=2, error bars are s.e.m.). Bar colours represent different cytokinin metabolite groups: free bases (yellow), ribosides (blue), glucosides (green). All protoplast samples derived from 9-day-old *Arabidopsis* seedling roots. See also Supplementary Fig. 1.

Although inactive cytokinin glucosyl-conjugates were relatively abundant in GFP+ cells, *trans-* zeatin (*tZ*) was the only active cytokinin significantly enriched in this population (Fig. 2a-Mock). In contrast, isopentenyl adenine (iP), which has similarly high affinities to cytokinin receptors^8,20,21^, was not enriched. The inferred leading role of tZ in cytokinin response was next tested using INCYDE (2-chloro-6-(3-methoxyanilino)purine) to block cytokinin degradation by CKX enzymes^22^, and resulted in both enhanced *TCSn::GFP* signal (Fig. 2b) and further elevation of tZ content in the *GFP*^+^ cells (Fig. 2a-INCYDE). The importance of tZ in root responses is also supported by genetic evidence showing severe root defects in *abcg14* mutants where *tZ* transport is impaired^23,24^.

**Fig. 2.**
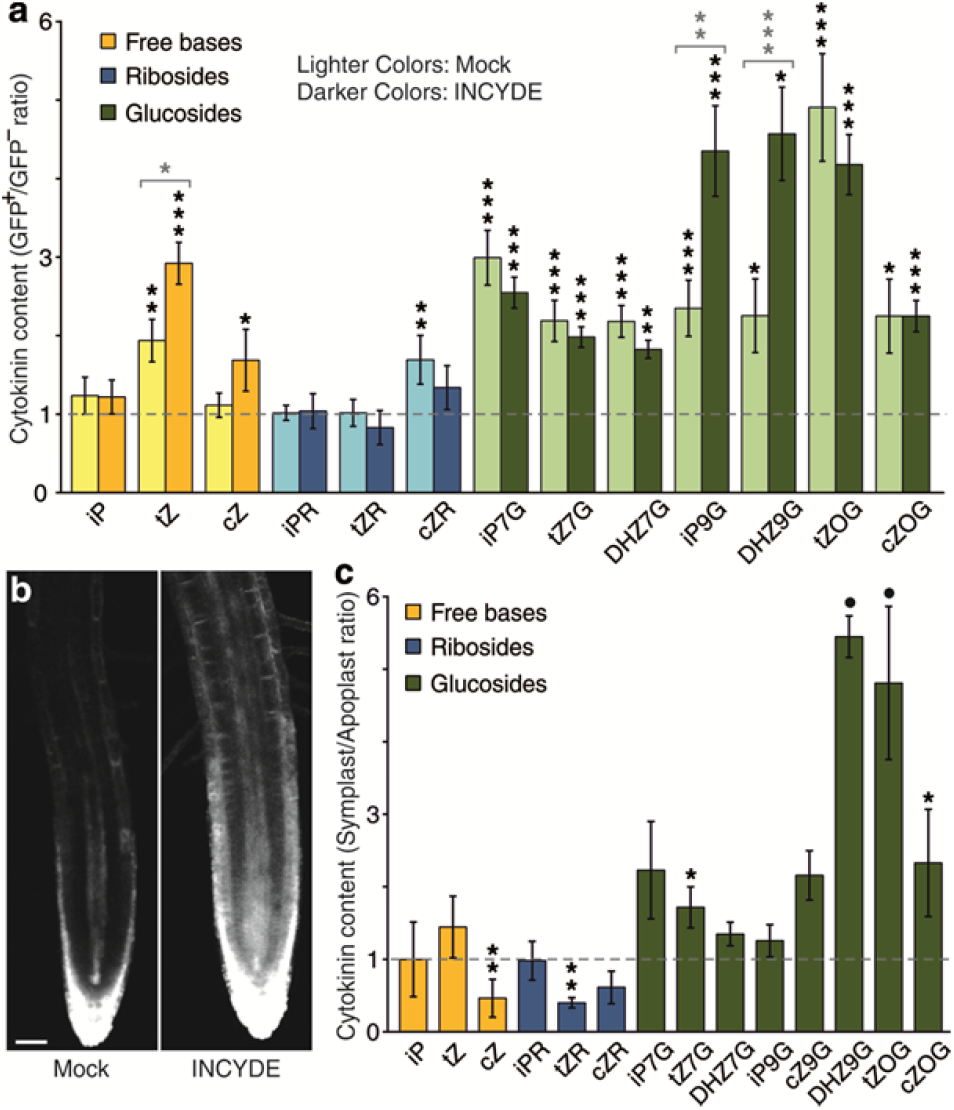
Cytokinin response is regulated by *trans-zeatin*. **a**, Ratio of cytokinin metabolite concentration between GFP^+^ and GFP^−^ protoplasts from 9-day-old *TCSn::GFP* seedlings roots, treated with or without 20 μM INCYDE during protoplasting. Cytokinins were quantified as fmol/100,000 protoplasts. Colours represent different cytokinin metabolite groups: free bases (yellow), ribosides (blue) and glucosides (green). Darker coloured bars are INCYDE-treated samples and lighter colours represent the corresponding mock treatment. Error bars are s.e.m. (n=9 for mock and n=6 for treated samples). Black stars indicate statistically significant differences in cytokinin concentration between GFP^+^ and GFP^−^ cells of *TCSn::GFP* treated or mock samples (Paired sample *t*-test). Grey stars above brackets denote statistically significant difference in cytokinin ratios between mock and treated experiment (one-way ANOVA and Tukey’s test). **b**, Confocal imaging of *TCSn::GFP* 5-day-old roots with or without 10 μM INCYDE treatment for 6 h. **c**, Ratios of cytokinin concentrations in symplastic/apoplastic fluid extracted from 9-day-old *Arabidopsis* wild-type roots. Ratios were derived from cytokinin levels calculated as fmol/g fresh weight of original tissue. n=3 pools of at least 1500 roots; error bars are s.e.m. Colour coding as in **a**. Stars indicate statistically significant differences in cytokinin concentration between symplast and apoplast by paired sample *t*-test. Significance levels are: dot (.), *P*<0.1; *, *P*<0.05; **, *P*<0.01; ***, *P*<0.001.

Since cell walls and extracellular space were absent from protoplast samples used in cell sorting (Fig. 2a), we additionally analysed apoplastic and symplastic fractions from roots, revealing relative enrichment of cytokinin glucosyl conjugates in the symplast (Fig. 2c), consistent with high levels detected in root protoplasts (Supplementary Fig. 2). However, these conjugate forms are essentially inactive (*26*) and unlikely to contribute directly to *TCSn::GFP* activation. Glucosyl-conjugate re-conversion to active forms during protoplast isolation was a possibility that was discounted by feeding labelled cytokinins (Supplementary Fig. 3). In contrast, cytokinin free bases and ribosides were either equally distributed between symplast and apoplast or enriched in the latter (Fig. 2c), opening up the possibility of extracellular cytokinins being able to initiate signalling.

We therefore next tested whether the bioactive cytokinins identified in the apoplast could be perceived by plasma membrane receptors^1,6,7,13,15,16,25^ by treating *TCSn::GFP* protoplasts with iP or *t*Z in free solution or covalently attached to Sepharose beads via flexible linkers designed to minimize steric hindrance to cytokinin binding (Supplementary Figs. 4 and 5a). Since the beads are much larger than the protoplasts (Supplementary Fig. 5a, red arrows), the attached cytokinin ligands were unable to enter the protoplast, and could thus be considered as membrane-impermeant signals (Fig. 3c). *TCSn* fluorescence signal strength after treatment with bead-bound cytokinins provided *in vivo* evidence for activation of cytokinin response through perception of extracellular cytokinins (Fig. 3a,b, also Supplementary Fig. 6b for *TCS::GFP* response). Further analysis on these samples showed that ~0.2% of the applied compounds had potentially been detached from the beads, corresponding to 4 nM free iP or *t*Z (Fig. 3c and Supplementary Fig. 7). Dose-response curves indicate that this concentration would not lead to a significant *TCSn::GFP* response (Fig. 3d), let alone the very large responses seen with both free and immobilised cytokinins. Notably, extracellularly restricted tZ resulted in a *TCSn::GFP* response approaching that elicited with 2 μM free *tZ*, whereas the response to immobilised iP was significantly lower (Fig. 3a,b). These results suggest that apoplastic tZ can trigger cytokinin response, consistent with the increase in *TCSn* signal when *tZ* degradation is impaired (Fig. 2a,b).

**Fig. 3.**
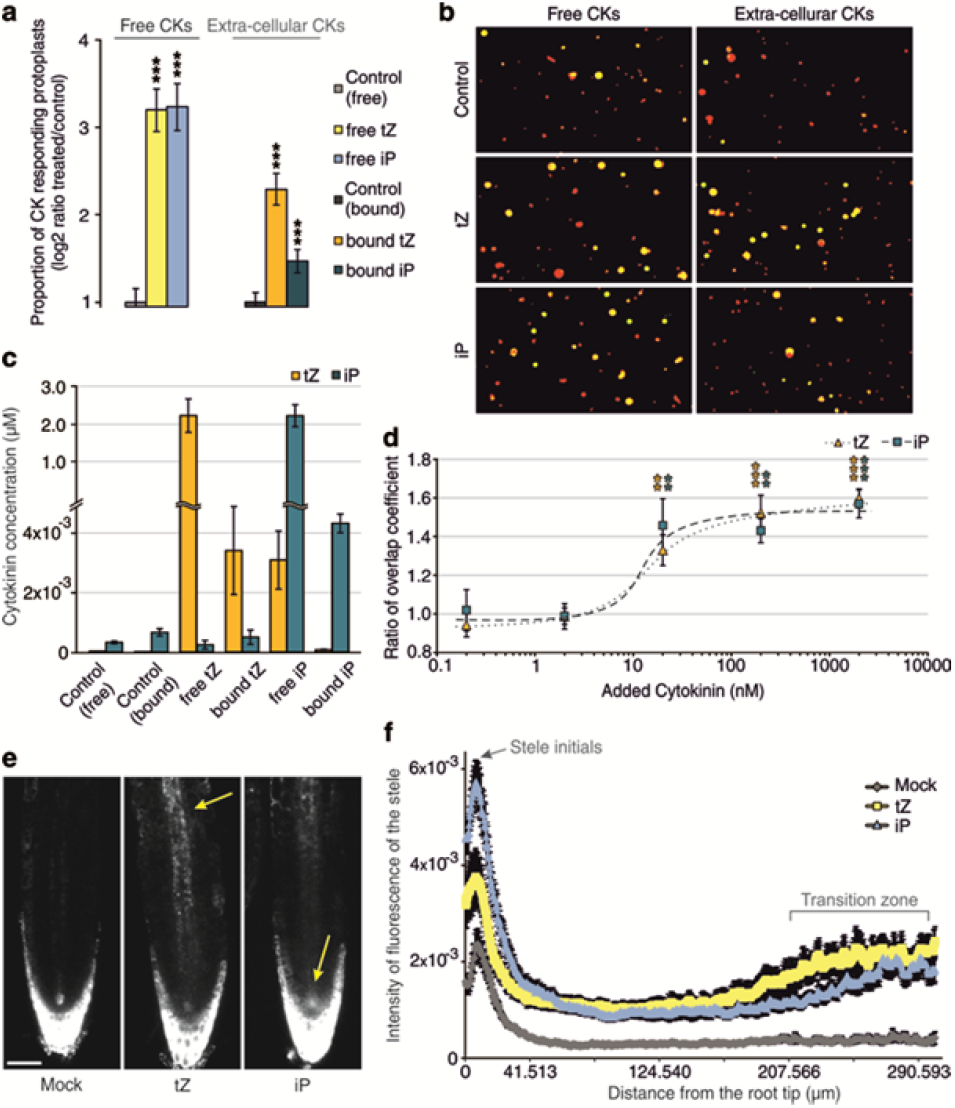
Extracellular cytokinins activate cytokinin response. **a**, Quantification of GFP fluorescence in protoplasts, derived from roots of 6-day-old *TCSn::GFP* seedlings, after treatment with or without free cytokinins (*t*Z or iP, 2 μM, denoted “free”) or immobilised cytokinins (*t*Z or iP, 2 μM equivalent, attached to Sepharose beads, denoted “bound”, also referred to as extracellular compounds). Negative controls without added cytokinin were incubations with and without beads (Control free and bound, respectively). ***, P<0.001 by one-way ANOVA and Tukey’s test, indicating significant differences in fluorescence intensity between control and corresponding free or extracellular cytokinin treatments. Three independent experiments were performed, with cytokinin treatment for 16 h then 1 μM FM4-64 applied 5 min prior to confocal imaging. Each data point represents n>20 images, corresponding to >1000 protoplasts; error bars are s.e.m. See also Supplementary Figures 4, 5 and 6b. **b**, Pictures of the treated protoplast samples described in **a** as indicated. **c**, Quantification of iP and *t*Z in the remaining protoplast samples from all three experiments described in **a. d**, Dose-response of GFP intensity in protoplasts of *TCSn::GFP* root protoplasts treated with 0.2, 2, 20, 200 or 2000 nM free *t*Z or IP for 16 h. Data are shown as ratio of treated/control. Error bars are s.e.m. and stars indicate significant differences in *TCSn::GFP* response by homoscedastic t-test (**, P<0.01; ***; P<0.001). Other details as for **a. e,f**, Responses of 6-day-old TCSn::GFP-expressing roots to treatment with 2 μM *t*Z or iP for 16 h. **e**, Confocal images of roots. Arrows indicate main zones of response to added cytokinins. **f**, Quantification of the GFP fluorescence intensity in the stele. Results are combined from two independent experiments, with the fluorescent signal of ten roots quantified in each experiment using ImageJ. Error bars are s.e.m.

Although iP, unlike *tZ*, was not enriched in the cytokinin-responsive cells (Fig. 2a) nor in the apoplast (Fig. 2c), exogenous supplies of both compounds triggered cytokinin responses in protoplasts (Fig. 3a,b). Cytokinin treatment of whole seedlings indicated differential spatial regulation of *TCSn* response by iP and *tZ*, as arrowed in Fig. 3e and annotated in Fig. 3f (also Supplementary Fig. 6a for corresponding *TCS::GFP* responses). iP had the strongest effect on meristematic stele cell initials, whereas *tZ* response was maximal in the transition zone in the stele. Although the exact biological role for cytokinins in the stele initials remains to be determined, *TCS* signal in those cells was absent in *ahk4* mutant roots^26^ (Supplementary Fig. 6c,d) providing genetic evidence for AHK4 being essential in cytokinin perception in the stele. We conclude that iP and tZ effects on cytokinin response and consequent biological functions can be tissue-specific, and may relate to their abundance or to differential sites of activity of particular cytokinin receptors. This is in accordance with previous findings showing greater tZ binding to receptors in maize roots compared with leaves^27^.

Finally, we tested the capacity of each of the three AHK receptors individually to respond to extracellular and free cytokinins, using *TCSn::GFP* lines mutated in the other two *AHKs*^26^. Although absolute signal strength was diminished in these mutant lines as shown elsewhere^26,28^, they all retained responsiveness to free IP and tZ (Fig. 4a,b). In root protoplasts, equivalent levels of response to both extracellular cytokinins was found for AHK2 (*ahk3 ahk4*), but interestingly AHK3 (*ahk2 ahk4*) and AHK4 (*ahk2 ahk3*) responded only to apoplastic IP or *tZ*, respectively. Application of IP and *tZ* to whole seedlings of the respective genotypes showed that AHK4 is not only essential but also sufficient for cytokinin response in the stele (Fig. 4b). In contrast, the same treatment of seedlings carrying only AHK3 or AHK2 receptors resulted in slightly enhanced *TCSn:GFP* response in some cell files within the stele (Fig. 4b arrows). Visualisation of AHK4-GFP and AHK3-GFP fusion protein by AiryScan super-resolution microscopy indicated that a proportion of both CK receptors was not co-localised with ER (Fig. 4c), similar to previous reports^4,5^ and to findings reported by Kubiasová et al (Supplementary File). Across multiple imaged cells, 24.6 ± s.e. 3.6% (n=3) of AHK3 and 35.5 ± s.e. 2.3% (n=5) of AHK4 signals were localised in non-ER regions. These imaging experiments further support our functional evidence showing that extracellular cytokinins can be perceived by the sub-populations of AHK receptor proteins that reside on the cell surface, and our data are consistent with experiments showing co-localisation of fluorescently tagged AHK4 and plasmamembrane markers (Supplementary file: Kubiasová et al.).

**Fig. 4.**
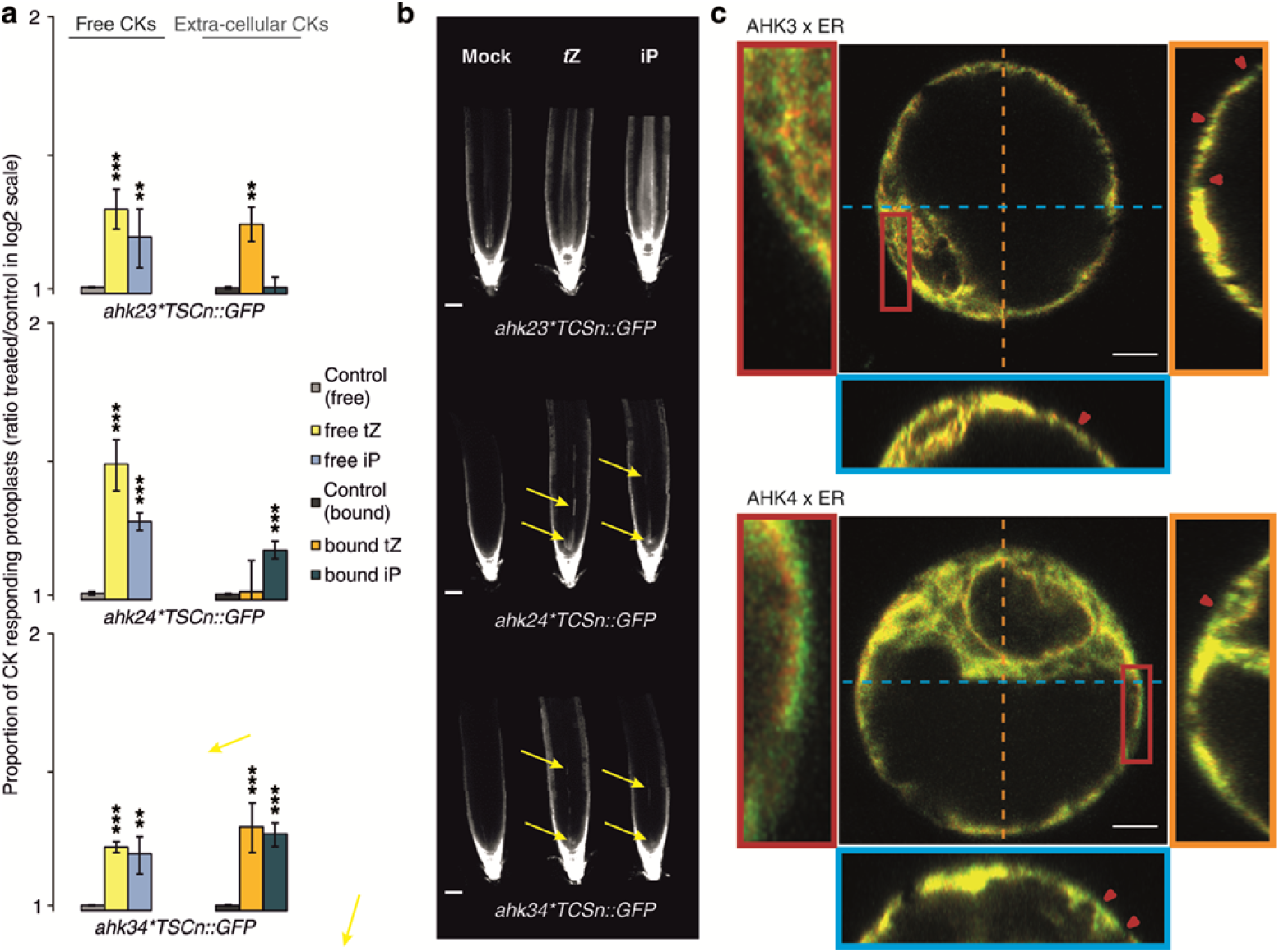
Cytokinin responses when only one AHK receptor is active, and receptor localisation. **a**, Quantification of GFP fluorescence in protoplasts derived from roots of 6-day-old *TCSn::GFP* seedlings in wild-type (Col-0), *ahk3,4, ahk2,3* and *ahk2,4* backgrounds, after treatment for 16 h with or without free cytokinins (*tZ* or iP, 2 μM, denoted “free”) or immobilised cytokinins (*tZ* or iP, 2 μM equivalent, attached to Sepharose beads, denoted “bound”). Negative controls without added cytokinin were incubations with and without beads (Control free and bound, respectively). **, P<0.01; ***; P<0.001 by one-way ANOVA and Tukey’s test, indicating significant differences in fluorescence intensity between control and corresponding free or extracellular cytokinin treatments. Three independent experiments were performed; each data point represents n>20 images, corresponding to >1000 protoplasts. Data are plotted on a log scale and error bars are s.e.m. See also Supplementary Figures 4 and 5. **b**, Confocal images of roots from *ahk3,4, ahk2,3* and *ahk2,4* double mutants expressing *TCSn:GFP*, after treatment for 24 h with 100 nM *tZ* or iP. **c,d.** Super-resolution 3D Airyscan images of Arabidopsis protoplasts expressing AHKs and the ER maker p24δ5. **c**, Example protoplast expressing AHK3-GFP (green) and p24δ5-RFP (red). The left panel shows a zoomed in region of the cell (red rectangle). The right panel depicts the YZ orthogonal view of the Z-stack (orange dashed line). The bottom panel shows the XZ orthogonal view of the Z-stack (blue dashed line). Red arrows indicate regions of AHK3-only signal on the cell surface. **d**, Example protoplast expressing AHK4-GFP and p24δ5-RFP. Other details as for **c**. Scale bars, 5 μm.

Several reports using AHK translational fusions to fluorescent proteins, point to AHK receptors predominantly located on ER membranes^3,4^ and show that AHK interaction with their downstream AHP partners can also occur at the ER^5^. However, plasma membrane localised receptors have never been excluded^4,27^ and AHK3 and AHK4 have been shown to at least partially reside in the plasma membrane^3,4^. Despite the strong evidence for ER location of AHKs, recent reviews have highlighted the lack of direct substantiation of extracellular cytokinin perception^1,6,7^. In this context, our multiple strands of evidence for responses to extracellular cytokinins initiated via plasma-membrane-bound receptors indicates that both sites of perception may exist.

In particular, our *in vivo* data point to perception of apoplastic *t*Z being an important route for cytokinin response activation in roots (Figs. 2c, 3a and 4a) while its symplastic degradation might act as a negative feedback loop in cytokinin signaling (Fig. 2a). These results are consistent with evidence showing that impaired cytokinin import/uptake results in induction of cytokinin response^25^ and with *t*Z specific binding by AHK4 in outer membranes^4^. Moreover, selective extracellular degradation by CKX expression leads to diminished cytokinin responses, whereas intracellular targeting of CKX did not have such an effect^25^. Nonetheless, intracellular inactivation of *t*Z by endogenous CKX does appear to occur (Fig. 2a-INCYDE;^29^).

The spatially distinct tissue-level responses to iP and *tZ* (Fig. 4a,b) may relate to ligand preferences^8,20,21^, sites of maximal expression^14^ of each AHK type, or/and differentially localised and expressed CKX enzymes^30^. The contrasting specific extracellular cytokinin responses in different AHK double mutant protoplast may reflect that the single remaining AHK in each case is only acting as a homodimer, whereas in WT plants heterodimerisation may occur^5^. Although localisation-related selectivity of receptor-molecule interactions merits further exploration, here we have demonstrated the presence of functional receptors at the plasma membrane of the cell, thus substantially resolving the lengthy debate on this issue^1,4–7,25,27^. We therefore propose that key regulatory elements leading to cytokinin response include selective molecule-receptor affinities and influences of tissue-specific apoplastic pH^31^, together with heterogeneous tissue- and cell-specific distribution of cytokinins, their cognate receptors and cytokinin inactivation enzymes. It remains to be ascertained whether different biological functions are associated with each location.

## SUPPLEMENTARY MATERIALS

**Supplementary Figures 1 to 7**

**Materials and Methods**

**References (37-42)**

## Author contributions

IA, CT and KL conceived the project; IA performed most of the experiments; IA, ON, LP and MK conducted the purification and quantification of cytokinins; IA, TV and SR discussed and performed the confocal experiments and developed semi-automated image processing for quantification; RS, VM and KD developed and produced the cytokinins attached to sepharose beads; MP and JH provided the homozygous lines of *ahk* mutant combinations with *TCSn::GFP* and *TCS::GFP;* IA and KO did the apoplastic fluid experiments; OP generated the *AHK::GFP* constructs and ZG, AJ and JF performed the respective transfection assays and protoplast imaging. IA; CT, and KL analysed and interpreted the data; IA and ON made the Figures; IA, CT and KL wrote the paper.

## Author Information

ORCID IDs: 0000-0001-9053-2788 (I.A.); 0000-0003-4938-0350 (K.D.); 0000-0002-2622-6046 (J.H.); 0000-0002-5603-706X (M.K.); 0000-0002-8153-907 (T.V.); 0000-0003-3452-0154 (O.N.); 0000-00025803-2879 (M.P.); 0000-0003-2537-4933 (L.P.); 0000-0002-0013-3239 (S.R.); 0000-0003-2901-189X (K.L.); 0000-0001-6635-1418 (C.T.); 0000-0003-0645-2019 (O.P.); 0000-0002-4481-6680 (R.S.); 00000003-4783-1752 (Z.G.); 0000-0002-8302-7596 (J.F.); 0000-0003-2901-189X (K.L.); 0000-0001-66351418 (C.T.);

## Acknowledgements

We thank Bruno Müller for critical discussions and provision of *TCSn::GFP* and *TCS::GFP* lines, and Roger Granbom (UPSC, Umeå Sweden) for technical assistance. The Swedish Metabolomics Centre and the IST Austria Bio-Imaging facility is acknowledged for access to instrumentation. The work was funded by the European Molecular Biology Organization (EMBO ASTF 297-2013) (IA), Development-The Company of Biologists (DEVTF2012) (IA; CT), Plant Fellows (the International Post doc Fellowship Programme in Plant Sciences, 267423) (IA; KL), the Swedish Research Council (621-2014-4514) (KL), UPSC Berzelii Center for Forest Biotechnology (Vinnova 2012-01560), Kempestiftelserna (JCK-2711) (KL). The Ministry of Education, Youth and Sports of the Czech Republic via the European Regional Development Fund-Project ‘Plants as a tool for sustainable global development’ (CZ.02.1.01/0.0/0.0/16_019/0000827) (ON, OP, RS, VM, LP, KD) and project CEITEC 2020 (LQ1601) (MP, JH) provided support, as did the Czech Science Foundation via projects GP14-30004P (MP) and 16-04184S (OP, KD, ON), Vetenskapsrådet and Vinnova (Verket för Innovationssystem) (TV, SR), Knut och Alice Wallenbergs Stiftelse via “Shapesystem” grant number 2012.0050. AJ was supported by the Austria Science Fund (FWF): I03630 to JF. The research leading to these results has received funding from European Union’s Horizon2020 program (ERC grant agreement n° 742985) and FWO-FWF joint project G0E5718N to JF. The authors declare that they have no conflicts of interest.

## Methods

### Plant material and growth conditions

The transgenic line *TCSn::GFP*^11^ was used for all cell-sorting experiments, *TCSn::GFP* and *TCS::GFP*^28^ were employed in protoplast and seedling treatment experiments, and the ecotype Col-0 was used for extraction of apoplast/symplast and for feeding experiments with labelled cytokinins. Crossed lines *ahk × TCSn::GFP* were published previously^26^ and the *ahk* mutant lines were crossed also with *TCS::GFP.* All experiments used seedlings grown on plates under axenic conditions. For cell sorting and confocal microscopy experiments, roots were harvested from 8 and 6 day old seedlings, respectively. Full details are in Supplementary methods

### Fluorescence activated cell sorting (FACS)

The distal half of 8 d old roots of *TCSn::GFP* seedlings were harvested, protoplasts were isolated in presence or absence of 20 μM INCYDE, and sorted using a BD FACS Aria I flow cytometer (BD Biosciences) as described^10^. Isolated protoplasts were kept at 4°C for 10 min and then loaded into the FACS. Healthy protoplasts were initially selected for size and granularity and then for autofluorescence and GFP fluorescence intensity. From loading into the FACS until reaching the fully isolated GFP^+^ and GFP^−^, or GFP^+^_max_ and GFP^+^_min_, protoplasts status (approximately 2 h), protoplasts remained at 4¼, then were immediately frozen in liquid nitrogen and stored at −80°C until cytokinin analysis. Each biological replicate represents an independent experiment.

### Protoplast feeding experiments

Protoplasts from roots of 8-day-old were resuspended in 2 ml of protoplast buffer (600 mM mannitol, 2 mM MgCl_2_, 10 mM KCl, 2 mM CaCl_2_, 2 mM MES, and 0.1% w/v BSA, pH 5.7). One μM of [^13^C_5_]trans-zeatin (*tZ*) or [^13^C_5_]cis-zeatin (cZ) was added to all protoplast samples except samples comprising the 0 min time-point which were harvested immediately; centrifuged at 1000 ×*g* for 3 min at 4°C, then frozen in liquid nitrogen and stored at −80°C until cytokinin analysis. After 30, 60 and 90 min of incubation in the dark at room temperature and under continuous shaking (46 rpm), treated samples were similarly collected. Cell numbers were estimated by hemocytometer.

### Isolation of apoplastic and symplastic fractions

Apoplastic and symplastic fluids were isolated from roots of 8-day-old Col-0 seedlings. Apoplastic fluid was collected by centrifugation at 900 *×g* for 20 min at 4°C. After freezing and thawing the remaining root tissue, symplastic fluid was collected by 15 min centrifugation at 2500 *×g* at 4°C. Presence of symplastic fluid in the apoplastic fractions was assessed to be 10 to 15% based on assays using the cytosolic malate dehydrogenase enzyme marker. Three biological replicates were analysed and each was a pool of at least 1500 seedlings. Full details are in Supplementary Methods.

### Cytokinin quantification

Cytokinins were purified using in-tip solid-phase microextraction and the cytokinin profile was quantitatively analysed by multiple reaction monitoring UHPLC-MS/MS (1290 Infinity Binary LC System coupled to a 6490 Triple Quad LC/MS System Agilent Technologies), as described^10^. Cytokinin concentrations were determined using Mass Hunter software (version B.05.02; Agilent Technologies) using stable isotope dilution. Labelled and endogenous cytokinin metabolites after protoplast treatment with^13^C_5_-*tZ* and^13^C_5_−*c*Z were also measured using LC-MS/MS, with the multiple reaction monitoring transitions as described^32^.

### Seedling treatments

Six-day-old *TCSn::GFP* or *TCS::GFP* seedlings were transferred liquid media with additions of 10 μM INCYDE, 2 μM *t*Z or 2 μM iP. After 6 h, samples treated with INCYDE were collected for confocal imaging, whereas samples treated with *t*Z and iP were similarly examined after 16 h. During incubation, samples were placed on an orbital shaker at 126 rpm under normal growth conditions. Seedlings of cytokinin receptor mutants (*ahk2,3, ahk3,4* and *ahk2,4* in *TCSn::GFP* background) were transferred for 24 h to solid media containing 100 μM IP or *tZ*.

### Generation of cytokinins attached to Sepharose beads

iP and *t*Z ligands possessing short linkers at the N^9^ position were synthesised according to the scheme in Supplementary Fig. 4, and confirmed by^1^H NMR spectra. Ligands were coupled to NHS-activated Sepharose^™^ 4 Fast Flow beads (GE Healthcare, United Kingdom). Control beads blocked with ethanolamine were prepared in the same way, omitting the ligand immobilisation step. Absorbance at 272 nm was used to determine the concentration of the immobilised cytokinin ligand. Full details are in Supplementary Methods

### Protoplast treatments

Protoplasts isolated from 6-day-old roots were resuspended in 4 mM MES (pH 5.7) buffer containing 0.5 M mannitol 20 mM KCl and 10 g/L sucrose. Protoplasts were mixed with free iP or *tZ* (2 μM), or with Sepharose beads with attached *tZ* or iP (2 μM equivalent) in wells of 96-well plates. Controls without added cytokinin were also examined with and without beads. Samples were incubated in darkness at room temperature under continuous shaking (46 rpm) for 16 h. Five min prior to confocal imaging 1 μM of FM4-64^33^ was added to stain all cells. Parallel aliquots were taken for cytokinin analysis to testing for possible leakage of cytokinins from the beads.

### AHK-GFP constructs for AiryScan imaging

The genomic sequences of the *AHK3* (At1g27320) and *CRE1/AHK4* (At2g01830) genes were inserted into pENTR-2B-Dual-AHK3 or pENTR-2B-Dual-AHK4 vectors, recombined into the p2GW7,0 vector containing the 35S promoter^34^, and the final constructs were used for protoplast transformations.

### AiryScan imaging and analysis

Protoplasts were derived from 4-d-old Arabidopsis root suspension culture and cotransformed with 35S::AHK-GFP construct and ER marker 35S::RFP-p24δ5^35^. Full details are in Supplementary Methods. After 12 h, protoplasts were transferred to 35 mm glass bottom MatTek dishes (coverslip thickness #1.5) coated with poly-L-lysine (Sigma) and imaged. Samples were imaged using a Zeiss LSM 880 inverted fast Airyscan microscope with a Plan-Apochromat 63x NA 1.4 oil immersion objective. Ten to 13.25 μm thick z-stacks of transformed cells were taken using Nyquist sampling steps. Images were then subjected to Airyscan processing. The channels were checked for correct alignment. The ER marker channel was then filtered with a Gaussian blur and converted to a mask in Fiji^36^. A custom Matlab script then determined the percentage signal present within and outside of the masked region in the channel of interest.

### Confocal microscopy

GFP expression was recorded using confocal laser scanning microscopy (Zeiss LSM780). The 488 nm laser line was employed for the GFP and FM4-64 fluorescence detection, and emission was detected between 490 and 580 nm and between 620 and 670 nm^33^, respectively. Full details are in Supplementary Methods.

## Reporting Summary

Further information on research design is available in the Nature Research Reporting Summary linked to this article.

## Additional information

**Supplementary information** is available for this paper at

**Correspondence and requests for materials** should be addressed to K.L. or C.T.

## Supplementary Information

**Supplementary Figure 1.**
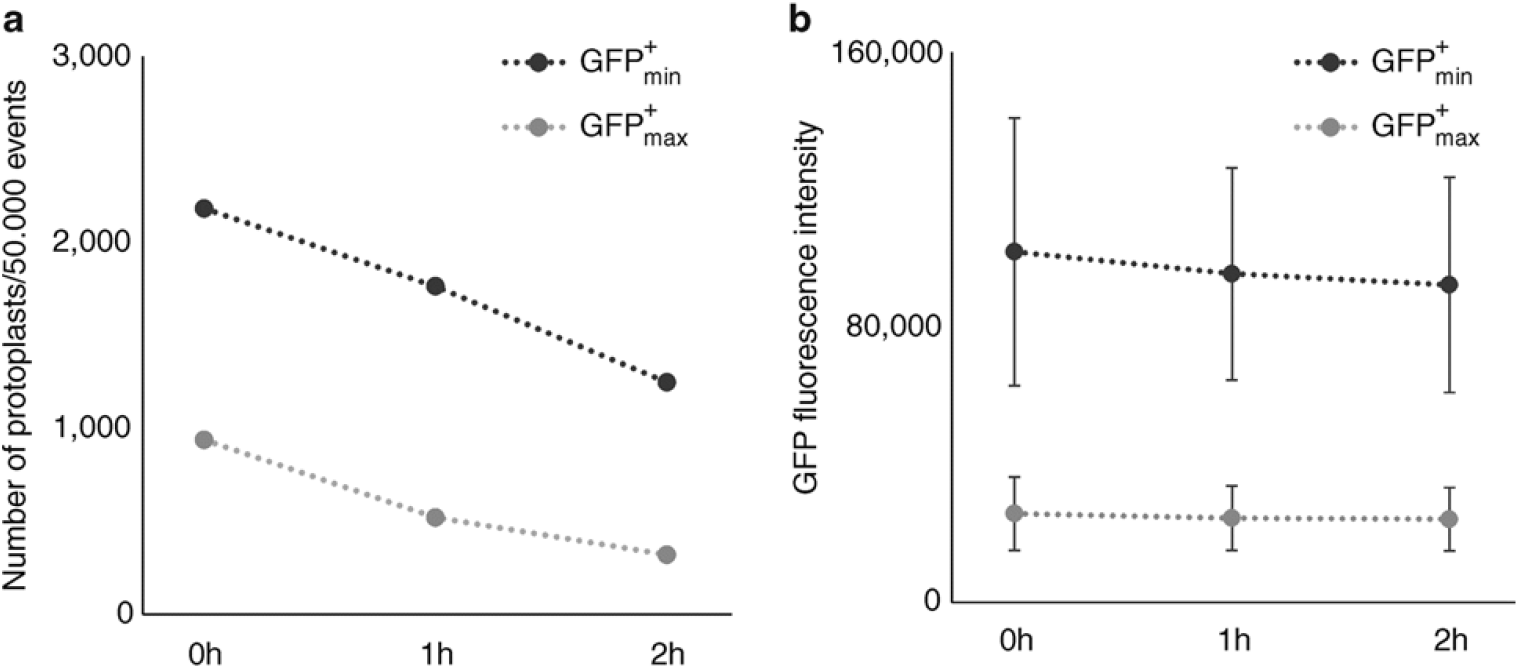
Control experiment testing GFP fluorescence stability over the 2 h of cell sorting. *TCSn::GFP* root protoplasts were sorted into GFP^+^ min and GFP^+^ max cell populations as shown in Fig. 1C. At 0, 1 and 2 h, 50.000 events were recorded and the GFP^+^ min and max protoplast populations were examined for **a**, their number and **b**, their GFP fluorescence. While there was a slight decrease in protoplast number this is consistent in GFP^+^_min_ and GFP^+^_max_ cell populations, indicating that there is no mixing between the two groups during sorting. The stability of GFP intensity during sorting and the respective significant difference between the two cell populations analysed are also shown.

**Supplementary Figure 2.**
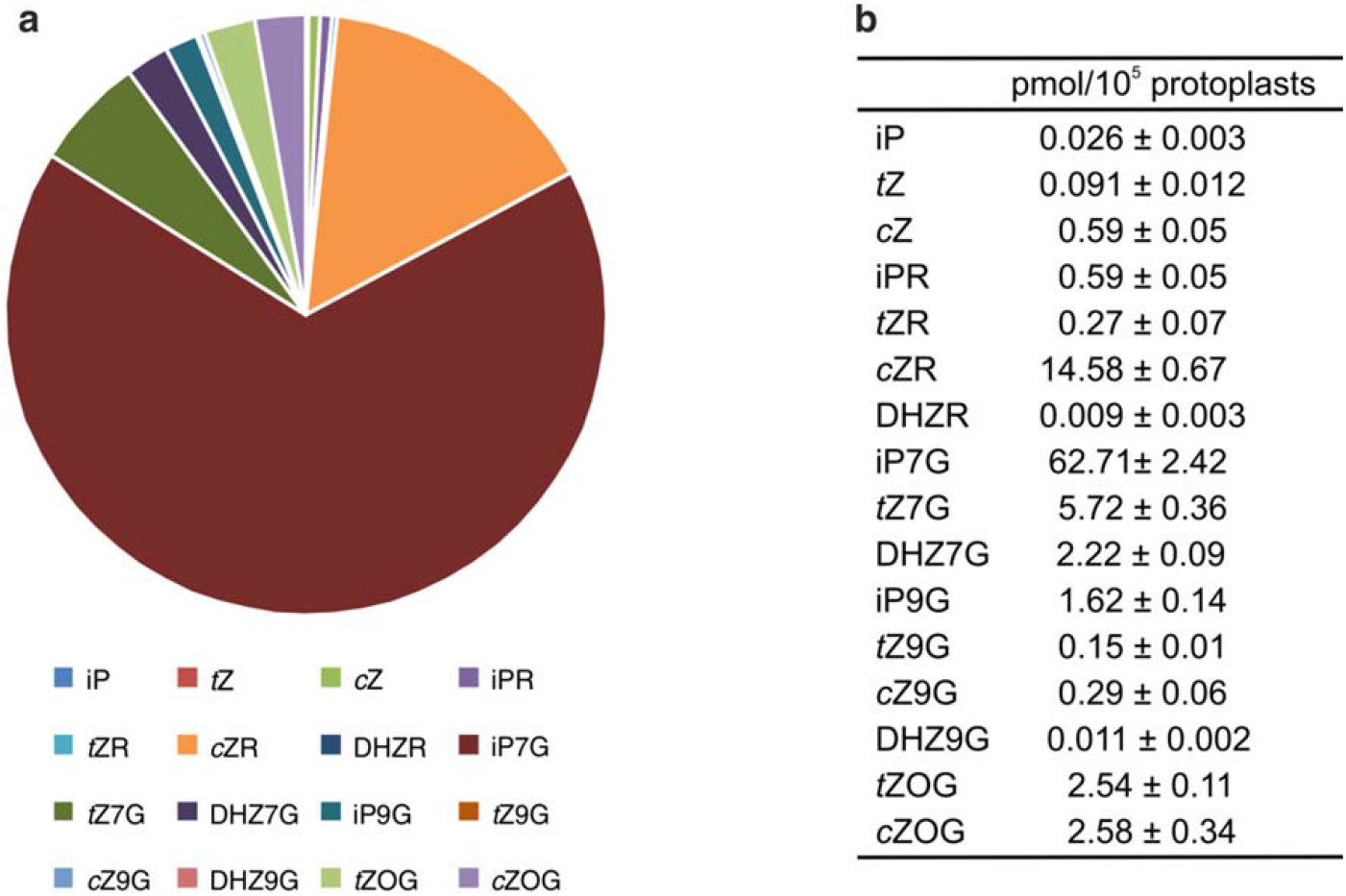
Cytokinin compound distribution and absolute levels in root protoplasts. Sum of cytokinin content in *TCSn::GFP*^+^ and *TCSn::GFP*^−^ root protoplasts, representing further processing of data in Fig. 2a-Mock. **a** Relative proportions of intracellular cytokinin compounds (% of total). **b** Absolute cytokinin quantification (fmol/100,000 protoplasts).

**Supplementary Figure 3.**
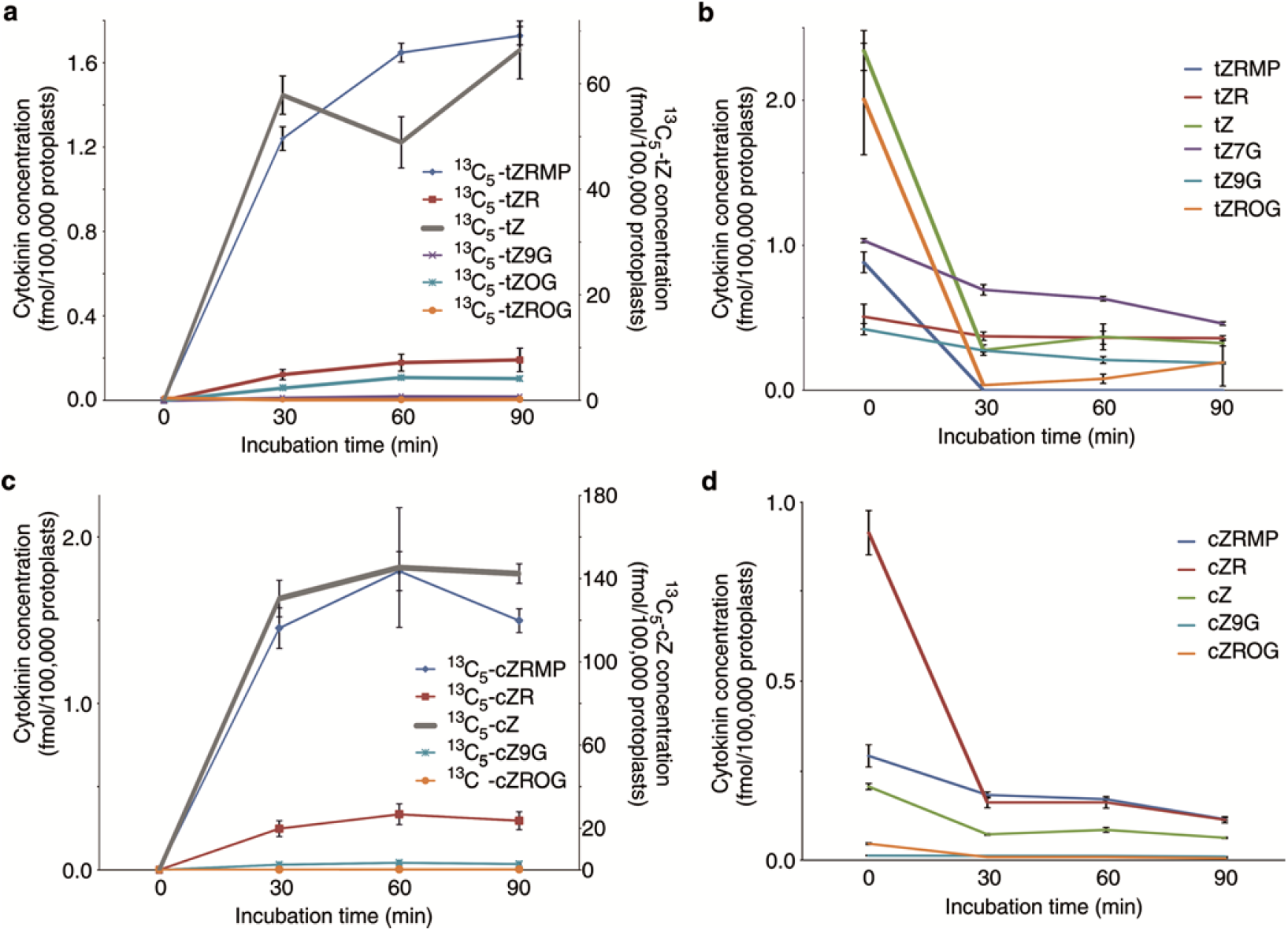
Uptake and metabolism of cytokinins by root protoplasts. Root protoplast suspensions supplied with 1 μM stable isotope-labelled^13^C_5_-*t*Z (**a** and **b**) or^13^C_5_-cZ (**c** and **d**). **a,c**, major labelled metabolites detected by LCMS. b,d, corresponding levels of endogenous (unlabelled) metabolites. Protoplast samples derived from 9-day-old Col-0 *Arabidopsis* seedling roots. Error bars represent s.e.m. While inactive precursor forms (^13^C_5_-*tZRMP* and^13^C5-*cZRMP*, respectively) were the main metabolites occurring in both treated protoplast samples (**a,c**), concentrations of most detected endogenous *t*Z- and *c*Z-cytokinins reduced over time, especially during the first 30 min after treatment (**b,d**). This suggests a tight homeostatic mechanism regulation on cytokinin metabolism with key components including the inactivation of active free bases (*t*Z and *c*Z) by conversion to cytokinin nucleotides, probably catalysed by APT enzymes.

**Supplementary Figure 4.**
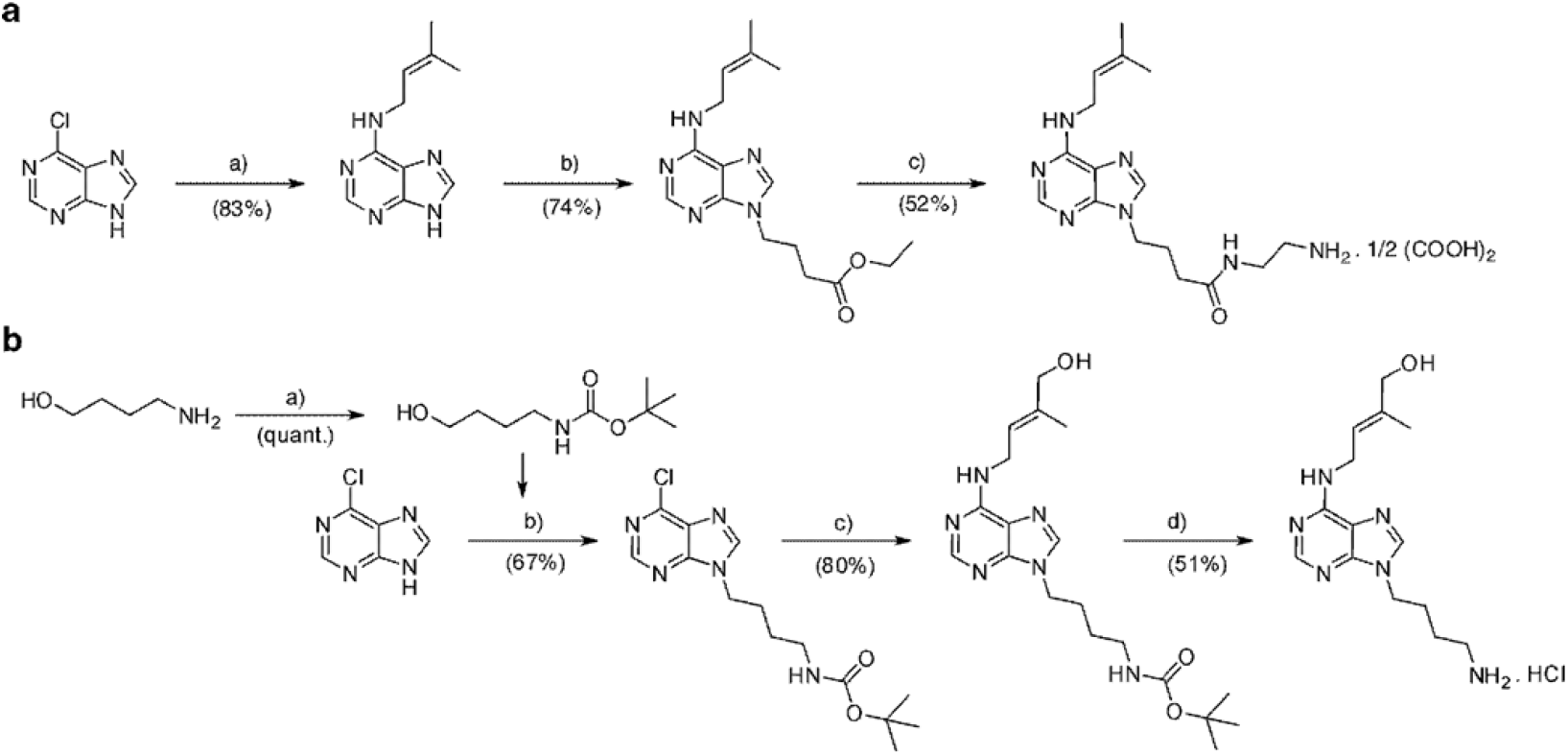
Reaction scheme for preparation of cytokinin ligands used for immobilisation on pre-activated chromatography beads. **a**, Synthesis of iP-type ligand: a) 1.2 eq. 3-methybut-2-en-1-amine hydrochloride, 2.5 eq. triethylamine, propanol, reflux, 4 h; b) 1.1 eq. ethyl 4-bromobutyrate, 2.5 eq. K_2_CO_3_, N,N-dimethylformamide, 16 h; c) 1/ 20 eq. ethane-1,2-diamine, reflux, 3 h; 2/ oxalic acid, methanol. **b**, Synthesis of tZ-type ligand: a) 1 eq. Boc anhydride, 3 eq. NaHCO_3_, tetrahydrofuran, methanol, 0°C then room temperature, 16 h; b) 1.1 eq. tert-butyl (4-hydroxybutyl)carbamate, 2 eq. triphenylphosphine, 2 eq. diisopropyl azodicarboxylate, tetrahydrofuran, 2 h; c) 1.3 eq. (E)-4-amino-2-methylbut-2-en-1-ol hemioxalate, 13 eq. triethylamine, methanol, 90°C, 6 h; d) 1/ Dowex 50W X8, dichloromethane, reflux, 6 h; 2/4 M methanolic NH3, 16 h; 3/ ethanolic HCl, methanol, 0°C. Reaction yields are shown in brackets.

**Supplementary Figure 5.**
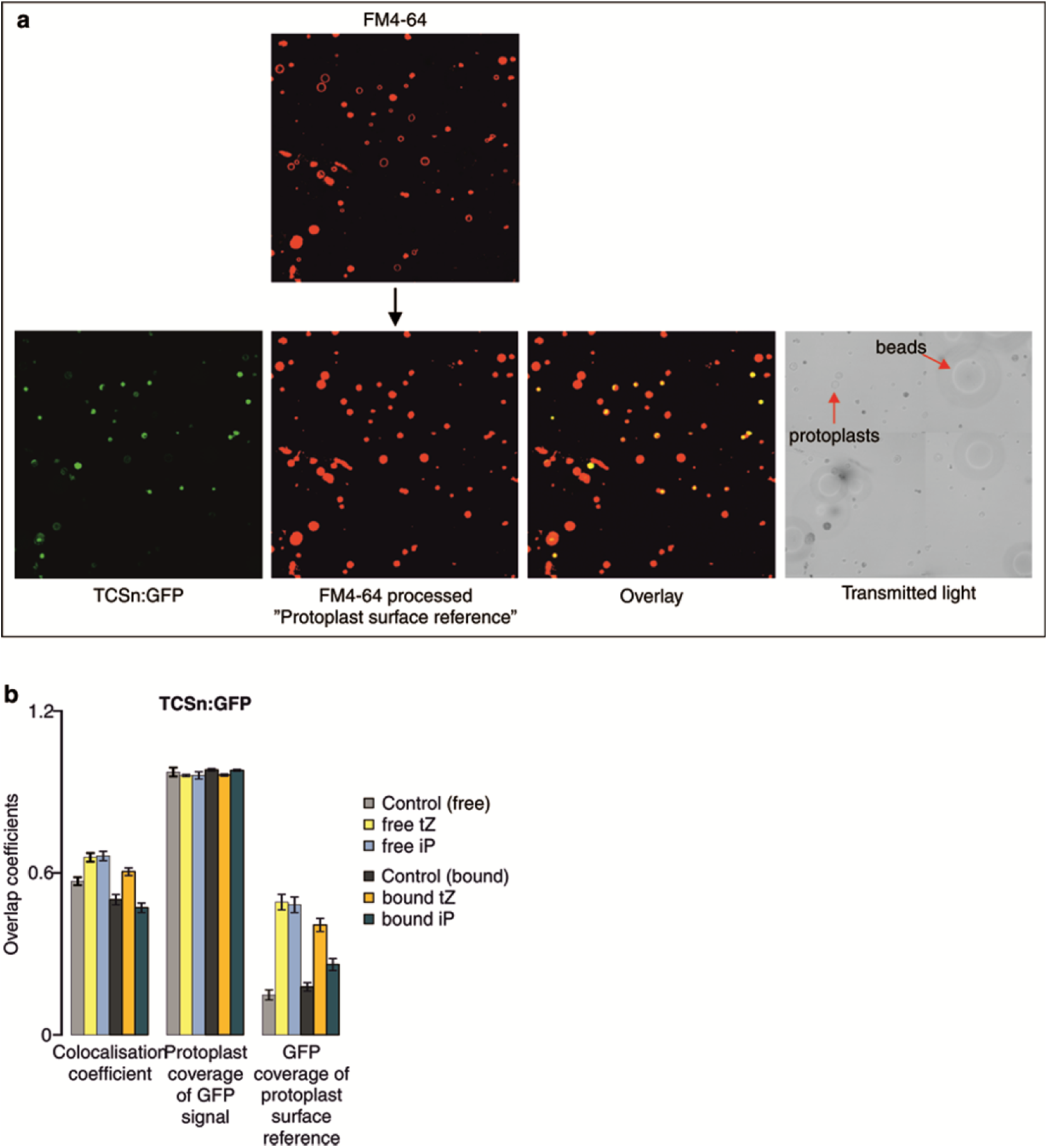
Quantification of GFP fluorescence intensity in root protoplasts. **a**, Illustration of image channels and overlay with indication of image processing method. **b**, Quantification results for *TCSn::GFP* protoplasts treated overnight (~16 h) with 2 μM free iP or *t*Z or with membrane impermeable iP or *t*Z (2 μM equivalent) linked to Sepharose beads (denoted “bound”, also referred to as extracellular compounds).

**Supplementary Figure 6.**
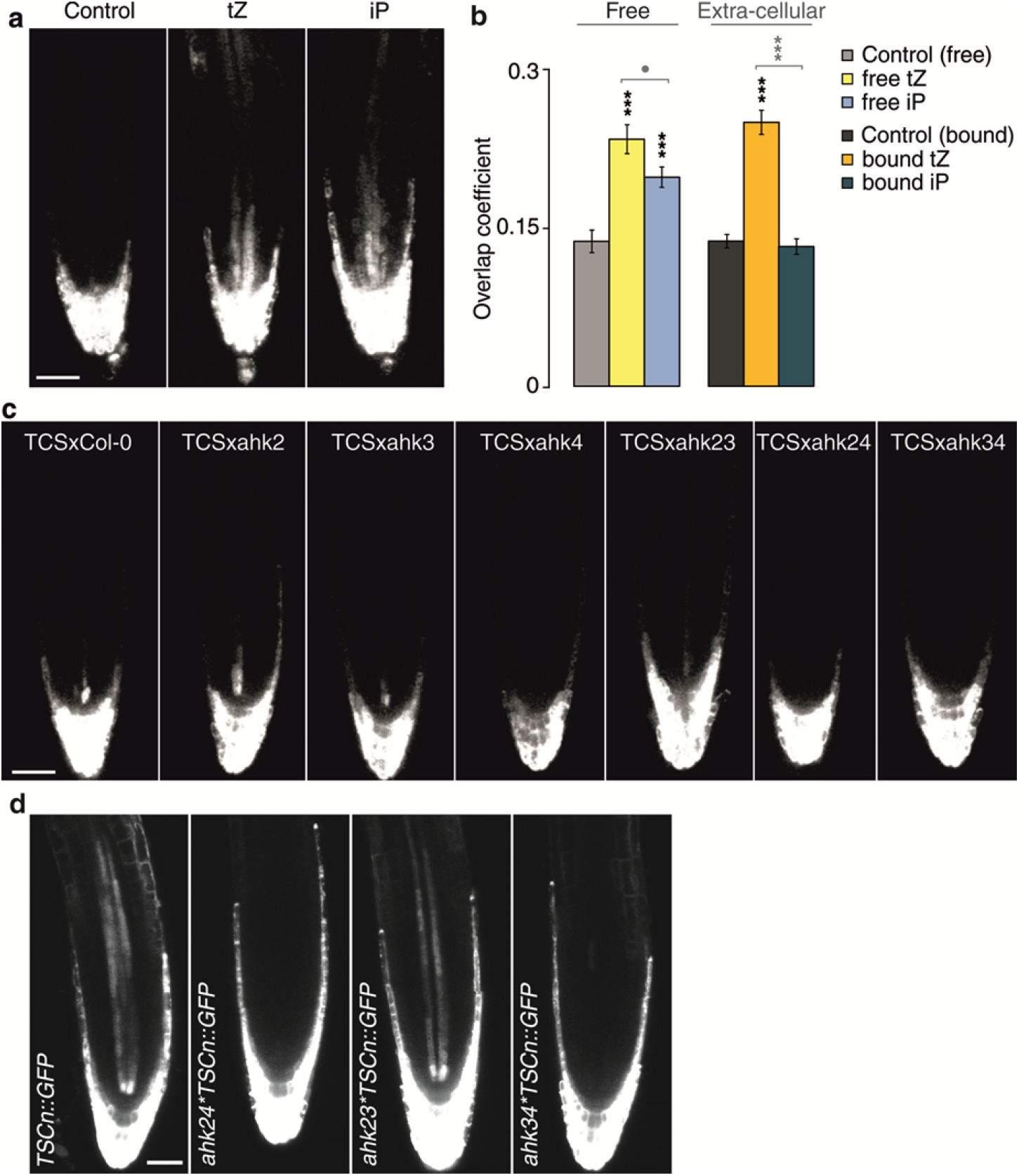
Cytokinin signalling responses to *tZ* and iP addition and to receptor mutations. **a**, *TCS::GFP* response 16 h after treatment with 2 μM iP or *tZ*. **b**, Quantification of GFP fluorescence in protoplasts, derived from roots of 6 day-old *TCS::GFP* seedlings, after treatment with or without free cytokinins (*tZ* or iP, 2 μM) or immobilised cytokinins (*tZ* or iP [2 μM] attached to Sepharose beads, denoted “bound”, also referred to as extracellular compounds). •, P <0.1, ***, P <0.001 by one-way ANOVA, and Tukey’s test indicating significant differences in fluorescence intensity between control and corresponding free or extracellular cytokinin treatments. Cytokinin treatment was applied for 16 h and 1 μM FM4-64 was added 10 min prior to confocal imaging. Each data point represents n>8 images, corresponding to >500 protoplasts. Cytokinin treatment of *TCS::GFP* seedlings (**a**) and protoplasts (**b**) resulted in similar response trends to *TCSn::GFP* in corresponding experiments (Fig. 3a,b). However, fluorescence intensities in *TCS::GFP* mock samples and in response to cytokinin treatments were lower than those observed with *TCSn:GFP.* **c**, TCS expression patterns in *ahk* mutant backgrounds. The *TCS::GFP* signal in stele cells is almost abolished when *AHK4* is mutated (*ahk4, ahk2,4, ahk3,4* mutants). **d**, Confocal images of 6-day-old roots comparing *TCSn::GFP* expression in wild-type (Col-0), *ahk2,4, ahk2,3* and *ahk3,4* backgrounds.

**Supplementary Figure 7.**
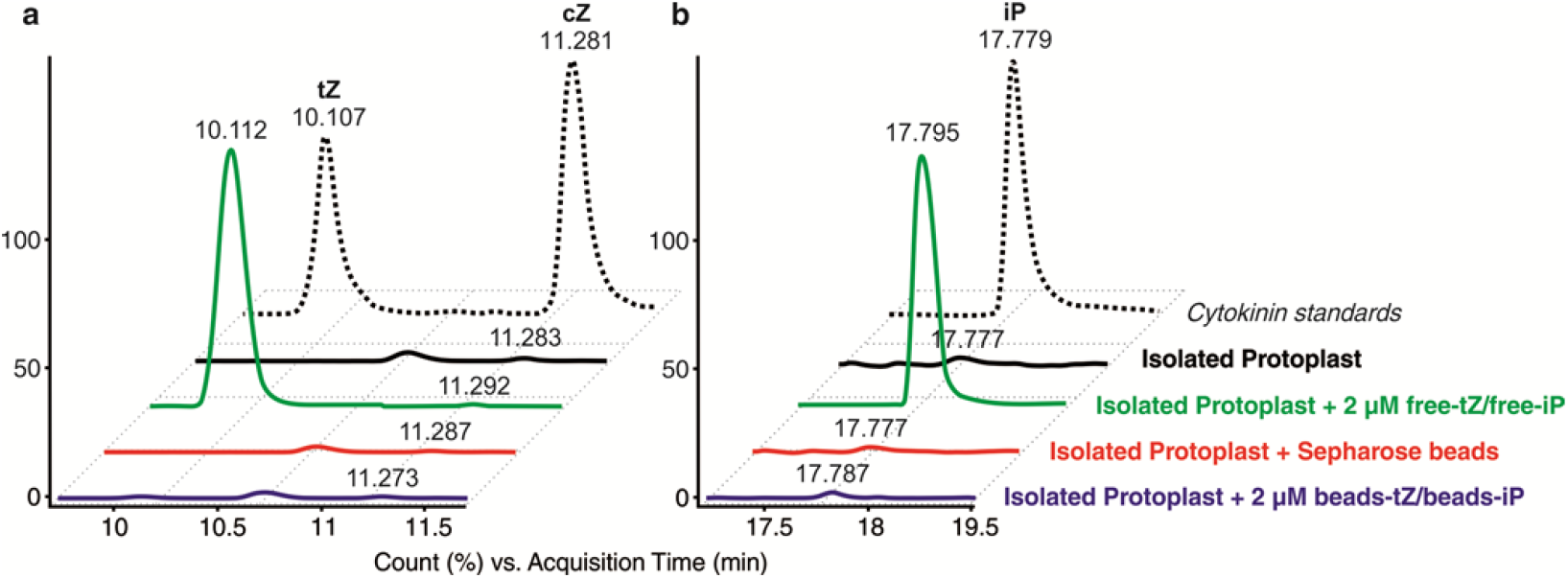
Sepharose-cytokinin beads stability tests. The possibility that some free *tZ* (**a**) or iP (**b**) might be released from *t*Z- and iP-Sepharose beads, respectively, was examined by LCMS analysis. The samples analysed were the same samples that were used for confocal imaging (Fig. 3a-c). The untreated protoplast profiles indicate low levels of endogenous cytokinins, whereas those treated with free cytokinins showed uptake of substantial excess of cytokinins. In contrast, incubation of protoplast with cytokinin beads resulted in minimal additional internal cytokinin.

## Supplementary Methods

### Plant material and growth conditions

All experiments used Arabidopsis (*Arabidopsis thaliana*). The transgenic line *TCSn::GFP*^11^ was used for all cell-sorting experiments, *TCSn::GFP* and *TCS::GFP*^28^ were employed in protoplast and seedling treatment experiments, and the ecotype Col-0 was used for extraction of apoplast/symplast and for feeding experiments with labelled cytokinins. Crossed lines *ahk/TCSn::GFP* were published previously^26^ and the mutant lines *cre1-2 (ahk4*) [14], *ahk2-5, ahk3-7, ahk2-5 ahk3-7 (ahk2,3), ahk2-5 cre1-2 (ahk2,4*) and *ahk3-7 cre1-2* (*ahk3,4*)^37^ were crossed also with *TCS::GFP.* Homozygous lines were then identified and used in confocal experiments. In all experiments, the seeds were surface sterilised with 20% (v/v) dilution of bleach for 5 min (2× 2.5 min) and then rinsed five times with sterile water. Seeds were sown in three rows (100 seeds/row) on square Petri dishes containing standard Murashige and Skoog medium (4.4 g/L Murashige and Skoog salt mixture, 1% sucrose, 0.5 g/L MES, 1% agar and adjusted to pH5.7 with KOH), covered with sterile mesh squares to facilitate the harvesting of the apical part of the primary root, and were stratified in darkness at 4°C for 3 d. Seedlings were then grown on plates placed vertically, for 8 d at 23°C under 150 μmol m^−2^ s^−1^ light with photoperiod of 16 h light and 8 h darkness. One standard cell-sorting experiment required 20 Petri dishes. For confocal microscopy experiments, sterilised seeds were sown (10 seeds/row), stratified and grown for 6 d as described above.

### Fluorescence activated cell sorting (FACS)

The distal half of 8 d old roots of *TCSn::GFP* seedlings were harvested, protoplasts were isolated in presence or absence of 20 μM INCYDE, and sorted using a BD FACS Aria I flow cytometer (BD Biosciences) as described^10,38^. The software used for data processing was BD FACSDiva version 6.1.2. Isolated protoplasts were kept at 4°C for 10 min and then loaded into the FACS. Healthy protoplasts were initially selected for their size and granularity and then for their autofluorescence and GFP fluorescence intensity. From loading into the FACS until reaching the fully isolated GFP^+^ and GFP^−^, or GFP^+^_max_ and GFP^+^_min_, protoplasts status (approximately 2 h), protoplasts remained at 4°C, then were immediately frozen in liquid nitrogen and stored at −80°C until cytokinin analysis. Each biological replicate represents an independent experiment.

### Protoplast feeding experiments

Protoplasts were isolated as described^10,38^ from roots of 8-day-old *Arabidopsis* seedlings and resuspended in 2 ml of protoplast buffer (600 mM mannitol, 2 mM MgCl_2_, 10 mM KCl, 2 mM CaCl_2_, 2 mM MES, and 0.1% w/v BSA, pH 5.7). One μM of [^13^C_5_]trans-zeatin (tZ) or [^13^C_5_]cis-zeatin (cZ) was added to all protoplast samples except samples comprising the 0 min time-point which were harvested immediately; centrifuged at 1000 ×*g* for 3 min at 4°C, then frozen in liquid nitrogen and stored at −80°C until cytokinin analysis. After 30, 60 and 90 min of incubation in the dark at room temperature and under continuous shaking (46 rpm), treated samples were similarly collected. Concentration of the samples was estimated by hemocytometer.

### Isolation of apoplastic and symplastic fractions

Apoplastic and symplastic sap were isolated from roots of 8-day-old Col-0 seedlings as described^39^ with modifications. The tissue was harvested, rapidly weighed and positioned in a 1 ml syringe without plunger. The syringe containing the sample roots was placed in a 25 ml centrifuge tube which was then centrifuged at 900×*g* for 20 min at 4°C. The syringe containing the remaining root tissue was frozen in liquid nitrogen then allowed to thaw at room temperature. Finally, the syringe was placed in a 25ml centrifuge tube into which the symplastic fluid was collected by 15 min centrifugation at 2500*×g* at 4°C. Collection of apoplastic fluid by this protocol has previously been shown to contain little cytoplasmic contamination^39,40^. Here, presence of symplastic fluid in the apoplastic fractions was assessed to be 10 to 15% based on assays using the cytosolic MDH enzyme marker. The different cellular origins of the two fractions were further confirmed by the resulting different cytokinin profiles of the “apoplast” and “symplast” shown in Fig. 2c. The cytokinin profile of the root symplast (Fig. 2c) was noted to be highly similar to the profile of the root protoplasts (Fig. 2a). Three biological replicates were analysed and each was a pool of at least 1500 seedlings.

### Cytokinin quantification

After frozen samples were thawed on ice, cytokinins were purified using in-tip solid-phase microextraction and the cytokinin profile was quantitatively analysed by multiple reaction monitoring UHPLC-MS/MS (1290 Infinity Binary LC System coupled to a 6490 Triple Quad LC/MS System with Jet Stream and Dual Ion Funnel technologies in positive ion mode; Agilent Technologies), as described^10^. Cytokinin concentrations were determined using Mass Hunter software (version B.05.02; Agilent Technologies) using stable isotope dilution. Labelled and endogenous cytokinin metabolites after protoplast treatment with^13^C_5_-*tZ* and^13^C_5_-*cZ* were also measured using LC-MS/MS, with the multiple reaction monitoring transitions as described^32^.

### Seedling treatments

Six-day-old *TCSn::GFP* or *TCS::GFP* seedlings were transferred into 6-well plates containing 2 ml of standard Murashige and Skoog liquid medium with additions of 10 μM INCYDE, 2 μM *tZ* or 2 μM iP. After 6 h, samples treated with INCYDE were collected for confocal imaging, whereas samples treated with *t*Z and iP were similarly examined after 16 h. During incubation, samples were placed on an orbital shaker at 126 rpm under normal growth conditions. Seedlings of cytokinin receptor mutants (*ahk2,3, ahk3,4* and *ahk2,4* in TCSn::GFP background) were transferred for 24 h to solid media containing 100 μM IP or tZ.

### Protoplast treatments

Protoplasts were isolated from 6-day-old roots of *TCSn::GFP, TCS::GFP, TCS::GFP ahk3* and *TCS::GFP ahk4* seedlings and resuspended in WI solution (4 mM MES (pH 5.7), 0.5 M mannitol and 20 mM KCl)^41^ supplemented with 10 g/L sucrose. 62.5 μL of protoplasts were transferred to wells in a 96-well plate and mixed with free iP or tZ (2 μM), or with Sepharose beads with attached *t*Z or iP (2 μM equivalent). Corresponding controls, without added cytokinin, were also examined with and without beads. Samples were incubated in darkness at room temperature under continuous shaking (46 rpm) for 16 h. Five min prior to confocal imaging 1 μM of the dye FM4-64^33^ was added to stain all cells. Twenty μL of the sample was used for imaging and the rest was frozen in liquid nitrogen and stored in −80°C until cytokinin analysis for testing possible leakage of cytokinins from the beads.

### AHK-GFP constructs for imaging

The genomic sequences of the *AHK3* (At1g27320) and *CRE1/AHK4* (At2g01830) genes were amplified from genomic DNA of Arabidopsis thaliana ecotype Columbia (Col-0) using the following primers: 5’-AATGTCGACGGATGAGTCTGTTCCATGTGC-3’ (forward) and 5’-ATTGCGGCCGCGATTCTGTATCTGAAGGCGAATTG-3’ (reverse) for *AHK3* and 5’-ATTGTCGACTGATGAGAAGAGATTTTGTGTATAATAATAATGC-3’ (forward) and 5’-ATTGCGGCCGCGACGAAGGTGAGATAGGATTAGG-3’ (reverse) for *CRE1/AHK4*, creating SalI linker sequences at the 5’ end, and NotI linker sequences at the 3’ end. The amplified PCR fragments were inserted into SalI and NotI sites of pENTR-2B-Dual (Invitrogen), yielding pENTR-2B-Dual-AHK3 and pENTR-2B-Dual-AHK4, respectively. Subsequently, enhanced GFP coding sequence was prepared using primers carrying NotI restriction site: 5’-AATGCGGCCGCACGGAGGTGGAGGTTCTATGGTGAGCAAGGGCGAGGAG-3’ (forward) and 5’-AATGCGGCCGCTTACTTGTACAGCTCGTCCATGCCG-3’ (reverse), and the resulting PCR fragment was inserted into the unique NotI site of pENTR-2B-Dual-AHK3 or pENTR-2B-Dual-AHK4 to obtain C-terminal GFP fusions with AHK3 and CRE1/AHK4, respectively. These entry clones were recombined using the Gateway LR reaction (Invitrogen) into the p2GW7,0 vector, containing the 35S promoter^42^, and the final constructs were used for protoplast transformations.

### Sample preparation for AiryScan imaging

Protoplasts were isolated from 4-d-old Arabidopsis root suspension culture in enzyme solution (1% cellulose; Yakult, 0.2% Macerozyme; Yakult in B5-0.34M Glc-mannitol solution; 2.2 g MS with vitamins, 15.25 g Glc, 15.25 g mannitol, H_2_O to 500 ml, pH to 5.5, with KOH) with slight shaking for 4 h, centrifuged at 1200 g for 5 min. The pellet was washed with B5-0.34 M Glc-mannitol solution followed by one time wash with B5-0.28 M sucrose buffer (0.44 g MS with vitamins, 9.6 g sucrose, H_2_O to 100 ml, pH to 5.5, with KOH) and resuspended in B5-0.34M Glc-mannitol solution to a final concentration of 2 × 10^5^ protoplasts per 50 μl. Protoplasts were cotransfected with 4 μg of 35S::AHK3-GFP or 35S::AHK4-GFP and with 4 μg of ER marker 35S::RFP-p24δ5 (45). DNAs were gently mixed together with 50 μl of protoplast suspension and 150 μl of PEG solution [0.1 M Ca(NO3)2, 0.45 M mannitol, 25% PEG 6000] and incubated in the dark for 30 min. PEG was washed by adding 0.275 M Ca(NO_3_)_2_ solution in two steps of 0.5 mL each, centrifuged at 800 *g* for 7 min and removed 240 mL of supernatant. The protoplast pellet was resuspended in 300mL of B5-0.34M Glc-mannitol solution and incubated for 12 h in the dark at room temperature. Protoplasts were transferred to 35 mm glass bottom MatTek dishes (coverslip thickness #1.5) coated with poly-L-lysine (Sigma) and imaged.

### Generation of cytokinins attached to Sepharose beads

iP and *tZ* ligands possessing short linkers at the N^9^ position were synthesised in the Department of Chemical Biology and Genetics, Centre of the Region Haná for Biotechnological and Agricultural Research, Czech Republic according to the scheme shown in Supplementary Fig. 4.^1^H NMR spectra were measured on a Bruker Avance 300 spectrometer at operating frequency 300.13 MHz in DMSO-d6 with chemical shifts (ppm) referenced to the solvent signal (residual [D5]DMSO = 2.50 ppm).^1^H NMR (300 MHz, DMSO-d_6_) δ (ppm), iP-type ligand: 1.67 (s, 3H,–CH3), 1.70 (s, 3H, –CH3), 1.97-2.07 (m, 4H, 2x –CH2-), 2.84 (t, J = 6.0 Hz, 2H, –CH2-), 3.26 (q, J = 6.0 Hz, 2H, –CH2-), 4.14-4.18 (m, 4H, 2x –CH2-), 5.31 (t, J = 5.4 Hz, 1H, -CH=), 5.35 (vbs, 2H, -NH2) 7.70 (bs, 1H, -NH-), 8.05 (t, J = 5.4 Hz, 1H, -NH-), 8.11 (s, 1H, -HAr), 8.19 (s, 1H, -HAr).^1^H NMR (300 MHz, DMSO-d_6_) δ (ppm), *t*Z-type ligand: 1.69 (s, 3H, -CH3), 3.26-3.42 (m, 2H, -CH2-), 3.82 (s, 2H, -CH2-), 4.27 (bs, 2H, -CH2), 4.58 (t, J = 5.8 Hz, 2H, -CH2-), 4.71 (bs, 1H, -OH), 5.57 (app. t, 1H, =CH-), 8.44 (bs, 4H, -NH3, -HAr), 8.60 (s, 1H, -HAr), 9.50 (bs, 1H, -NH-). NHS-activated Sepharose^™^ 4 Fast Flow beads (GE Healthcare, United Kingdom) were washed with a threefold volume of anhydrous DMSO and resuspended in equal volume of 20 mM solution of cytokinin ligand in DMSO. Coupling was performed at room temperature under continuous moderate stirring overnight. Subsequently the beads were washed in three successive steps with DMSO, DMSO:water (1:1, v/v) and water, and remaining non-reacted N-hydroxysuccinimide groups were inactivated with 1 M ethanolamine according to the manufacturer’s instructions. The cytokinin-coupled beads were rinsed then with water and ethanol:water (1:4, v/v) and stored in the latter solution at 4°C until required. Control beads blocked with ethanolamine were prepared in the same way, omitting the ligand immobilisation step. To determine the efficiency of coupling reaction, an aliquot of the cytokinin-coupled beads was thoroughly resuspended in glycerol:water (1:1, v/v) and its absorbance was measured at 272 nm. From this value, the absorbance of the blank beads was subtracted and the concentration of the immobilised cytokinin ligand was calculated following Lambert-Beer’s law.

### Confocal microscopy

GFP expression patterns in 6 day-old seedlings or isolated protoplasts of the transgenic *Arabidopsis* lines *Col-0, ahk3, ahk4, ahk2, ahk2,3, ahk2,4 and ahk3,4* carrying *TCSn::GFP, TCS::GFP, AHK3::GFP* or *AHK4::GFP* were recorded using confocal laser scanning microscopy (Zeiss LSM780). The 488 nm laser line was employed for the GFP and FM4-64 fluorescence detection, and emission was detected between 490 and 580 nm and between 620 and 670 nm^33^, respectively. Two tile scans were performed for root imaging and 5×5 tile scans for protoplast imaging. ImageJ (http://imagej.nih.gov/ij/) was used to quantify GFP fluorescence intensity. Fluorescence profiles of the stele and the full root were extracted using the Plot profile function. Plot profiles represent quantification from 10 roots per treatment and two independent experiments were performed. For the calculation of overlap coefficients (Figs. 3a and 4a and Supplementary Fig. 5b) in treated protoplasts, semi-automated digital processing was performed using ImageJ. Raw images have been converted into 8-bit. Noise was reduced using a Median filter. FM4-64 channel was converted to binary and Fill holes function was used to obtain the surface of the protoplasts as filled circular structures. Protoplasts were counted and extracted from cellular debris using surface and circularity thresholds using the Analyze particle function. Quantification of the *GFP* fluorescence in protoplasts was performed using the ImageJ plugin JACoP. Colocalisation coefficient corresponds to the Pearson’s correlation coefficient between the two channels (FM4-64 and GFP). The GFP signal overlapping the protoplast reference surface was measured using the Manders’ overlap coefficient (M2) and as an additional control on the reference values, the coverage of the protoplast on the GFP signal was also measured (M1) (Supplementary Fig. 5).

